# Infection dynamics in *L.major* autophagic machinery is regulated by ATG9-PI3P negative feedback loop

**DOI:** 10.1101/398545

**Authors:** Dipali Kosey, Bhavnita Soni, Prajakta Nimsarkar, Shailza Singh

## Abstract

Autophagy is a self-destructing mechanism of cell via lysosomal degradation, which helps to degrade/ destroy hazardous substances, proteins, degenerating organelles and recycling nutrient. It plays an important role in cellular homeostasis and regulates internal environment of cell, moreover, when needed causes non-apoptotic programmed death of cell. It’s usually detailed with ageing and nutritional stress, but recent researches provide support for its role in innate as well as adaptive immunity. Autophagy has been observed as one of the major factors in parasite clearance in leishmaniasis. Due to intra-cellular pathogen, the cell mediated response is only alternative for adaptive immunity against *Leishmania* in host. T-cells have been observed as main component of cell mediated immunity in leishmaniasis. The differentiation of T-cells generates either destruction or proliferation of parasite as different kind of cytokine mediate different results. Pro-inflammatory cytokine IL12 and TNFα generate Th2 response which helps in active phagocytosis of parasite whereas an anti-inflammatory cytokine like IL10 mediate parasite promotion by blocking autophagic pathways and inhibiting phagocytic actions. TLR2/6 mediated signaling stimulated by LPG produces many pro-inflammatory cytokines like IL12, TNFα and IL6 etc. In macrophages, it is found that TNFα via autocrine signaling induce autophagy and help in parasite killing of *Toxoplasma gondii* via JNK phosphorylation in macrophages. The phosphorylated JNK induces Beclin by phosphorylating Bcl2, which inhibits Beclin action, thus releasing Beclin and promoting VpS34 which recruits PI3P for autophagosome formation. Recruitment of PI3P is controlled by ATG9 protein, autophagy related gene protein, which in normal condition is inhibited by mTOR via ULK/ ATG1. IL10 has been observed to inhibit autophagy via starvation induced AKT-PI3K pathway in macrophages and result in inducing mTOR. Thus, we can conclude that IL10 inhibits recruitment of PI3P via mTOR. Through systems aspect, here, we decipher that Atg9-PI3P acts as a negative feedback loop in autophagic machinery of leishmaniasis.

## 1. Introduction

Leishmaniasis is the second most neglected disease after malaria, caused by obligatory intracellular protozoan parasite from genus *Leishmania.* The parasite has its digenetic life cycle completed between phlebotomine sandfly and mammalian host. During its life cycle in mammalian host it remains in its promastigote form and infect Natural killer (NK) cells where it survives, making NK cells as Trojan horse for infecting other major immune cell such as macrophage and dendritic cells^1^. Through its autophagic machinery the parasite convert itself into more virulent amastigote form^2^. The parasite targets the protector cells of immune system making them impaired results in disease development. The present work focuses on exploring autophagic machinery of host and the parasite through systems biology approach to find out important targets that will be exploited in future to develop therapeutics for leishmaniasis.

### 1.1 Immunobiology of Leishmaniasis

Toll-like receptors and the associated signalling pathways represent the most ancient host defence mechanism found in almost all eukaryotes from insects to mammals. The TLRs are 11 membered family (TLR1 to TLR11) having specificity for different pathogens/ pathogenic components and induce production of different cytokines. Leishmania parasite through Lipophosphoglycan (LPG) interacts with Toll like receptor present on macrophages specifically TLR2 and TLR6 and initiates MyD88 dependent TLR signalling^3,4^.

As the parasite is intracellular in nature, IgG not only fails to provide protection against this pathogen, but it actually contributes to disease progression. Thus, humoral branch of immunity does not play any role in parasite clearance^4^. The differentiations of T-cells generate either destruction or proliferation of parasite as different kind of cytokine mediate different results. Pro-inflammatory cytokine generate Th1 response, which helps in active phagocytosis and oxidative burst whereas an anti-inflammatory cytokine mediate parasite promotion by blocking autophagic pathways and inhibiting phagocytic actions as well as blocking nitric oxide production of host cell. The clearance or infection persistence depends on macrophage differentiation into M1 macrophage or M2 macrophage, which in turn depends on cytokine induction. M1 macrophages are induced by pro-inflammatory cytokines from initial M0 phase, which induce Th1 response of T-helper cells are activated for parasite clearance and the vice versa case is seen when M0 macrophages are induced by anti-inflammatory cytokines like IL10 to undergo changes and differentiated as M2 macrophages which encourage Th2 responses, thus prevailing the manifestation of parasite^4^.

### 1.2 Autophagy and Leishmaniasis

Autophagy is a self-destructing mechanism of cell via lysosomal degradation, which helps to degrade/ destroy hazardous substances, proteins, degenerating organelles and recycling nutrients. It plays an important role is cellular homeostasis and regulates internal environment of cell when needed causes non-apoptotic programmed death of cell. It usually was detailed with ageing and nutritional stress, but recent researches provide support for its role in innate as well as adaptive immunity^5^. It is further classified into mitophagy, chaperone mediated autophagy and macrophagy^6^. Macrophagy is found to be related to immunity of host^5^.

#### 1.2.1 Autophagy-Mechanism

##### Induction and autophagophore formation

The molecular machinery of autophagic processes involves several conserved Atg proteins called Autophagy-related-gene products. Various stimuli including nutrient starvation, elimination of growth factors etc., lead to the formation of the autophagophore. This step involves two protein complexes: A complex that contains the class III PI3K/Vps34, Atg6/Beclin1, Atg14 and Vps15/30 and a complex that includes the serine/threonine kinase Atg1/ULK1 (human homologue of Atg1 of yeast), a very important as well as an essential positive regulator of autophagosome formation^6^.

##### Phagophore elongation and autophagosome formation

The characteristic double-membrane autophagosome is formed in elongation step. It comprises two ubiquitin-like conjugation pathways, both catalysed by Atg7. The first system leads to the conjugation of Atg5-Atg12, which later forms a multimeric complex with Atg16L. The Atg5-Atg12-Atg16L complex associates with the outer membrane of the extending phagophore and engages for further process.

The second system marks in the processing of LC3, (mammalian homologue of the yeast Atg8). Upon induction LC3B is proteolytically sliced by Atg4 to generate LC3B-I. This molecule is activated by Atg7 and later conjugated to phosphatidylethanolamine (PE) in the membrane to produce processed LC3B-II. Processed LC3B-II is recruited onto the growing autophagophore and its integration is dependent on Atg5-Atg12^6^.

Unlike Atg5-Atg12-Atg16L, LC3B-II is found on both the inner and outer surfaces of the autophagosome, where it is required for the expansion and completion of the autophagic membrane. After closure of the autophagosomal membrane, the Atg16-Atg5-Atg12 complex dissociates from the vesicle, whereas a portion of LC3B-II remains covalently bound to the membrane (Fig 1). Therefore, LC3B-II can be used as a marker to monitor the level of autophagy in cells^7^. Recent studies validate self-multimerization and phosphorylation of Atg9 aid membrane tethering and/or fusion^8^.

**Fig. 1:**
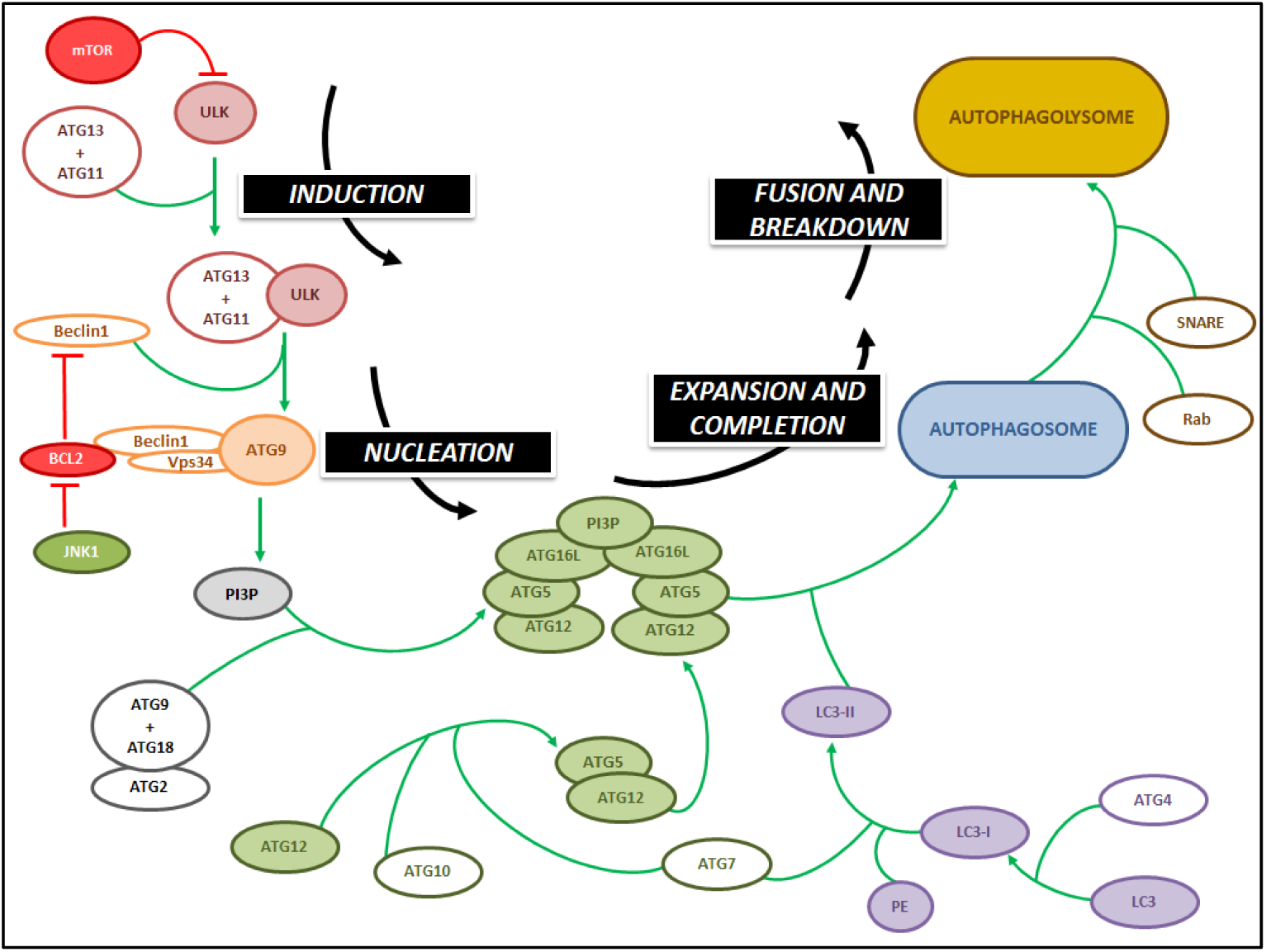
Molecular mechanism of autophagy.

**Fig. 2:**
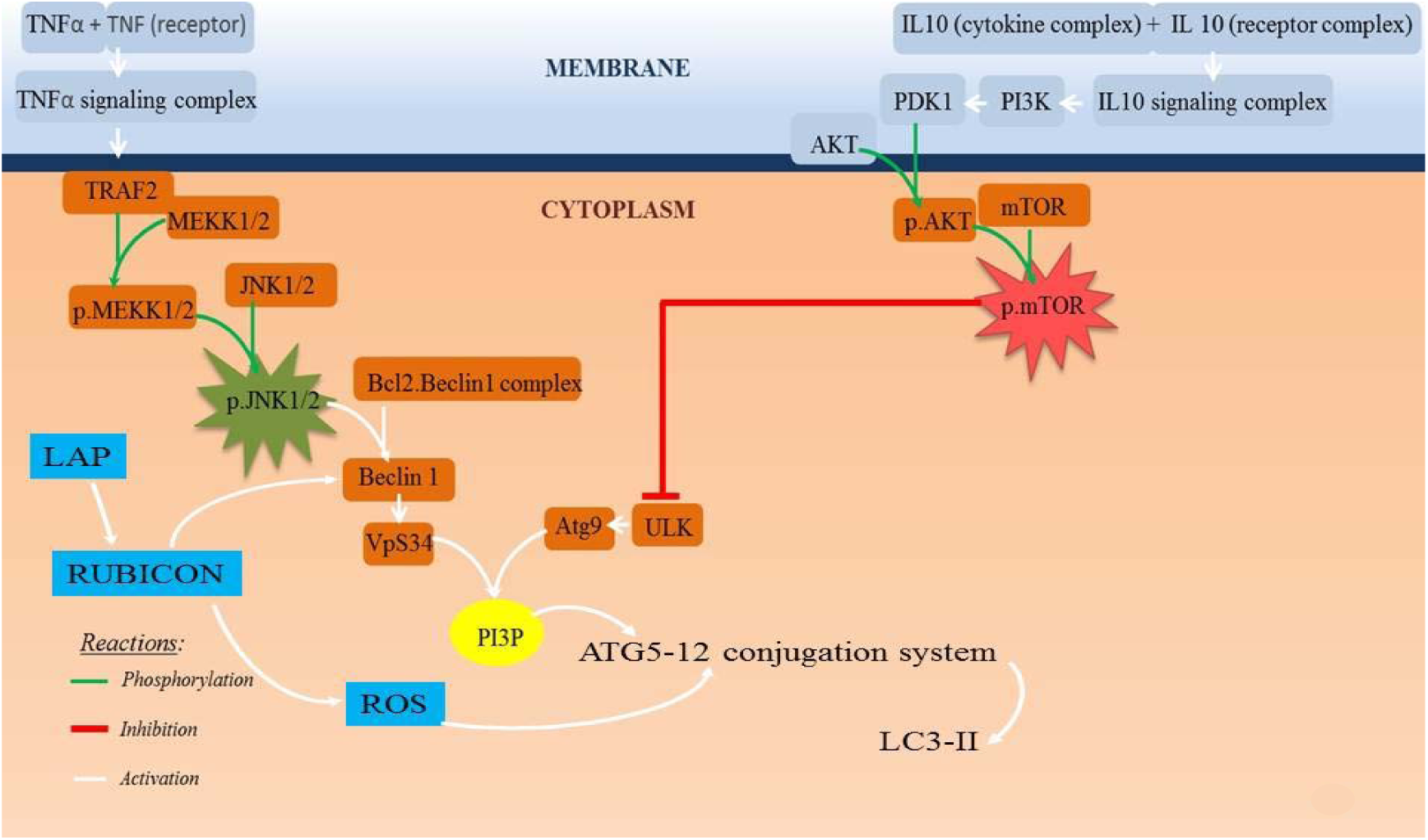
Signaling pathways connecting canonical and non-canonical autophagy.

##### Fusion, degradation and recycling

When the autophagosome formation is accomplished, LC3B-II attached to the outer membrane is cut from PE by Atg4 and released back to the cytoplasm. The fusion of the autophagosome and the lysosome in thought to require the lysosomal membrane protein LAMP-2 and the small GTPase Rab7.

After fusion, a series of acid hydrolases are involved in the degradation of the confiscated cytoplasmic cargo. The small molecules resulting from the degradation, particularly amino acids are transported back to the cytosol for protein synthesis and maintenance of cellular functions under starvation conditions^6^.

#### 1.3 Autophagy in Leishmaniasis

Autophagy has been observed as one of the major factors in parasite clearance in leishmaniasis^9^. Due to intra-cellular pathogen, the cell-mediated response is only alternative for adaptive immunity as discussed before via TLR signaling.

##### Interplay between TNFα and IL10 for autophagy in macrophages

TLR2/6 mediated signalling stimulated by LPG produces many pro-inflammatory cytokines like IL12, TNFα and IL6 etc4. In macrophages, it is found that TNFα via autocrine signalling induce autophagy and help in parasite killing of Toxoplasma gondii via JNK phosphorylation in macrophages^10^. The phosphorylated JNK induces Beclin by phosphorylating Bcl2, which inhibits Beclin1 action, thus releasing Beclin1 and promoting VpS34 that recruits PI3P for autophagosome formation. Recruitment of PI3P is controlled by ATG9 protein, autophagy related gene protein, which in normal condition is inhibited by mTOR via ULK/ ATG1 (Fig.2).

IL10 has been observed to inhibit autophagy via starvation induced AKT-PI3K pathway in macrophages via inducing mTOR^11^. Thus, we can conclude that IL10, which is found to be encouraged by parasite interaction to host, obstructs recruitment of PI3P via mTOR and eventually checks autophagy for parasite survival.

#### 1.3.2 Interaction between canonical and non-canonical autophagy

For the action of internalisation of parasite, *Leishmania* evades macrophages by phagocytosis but recruitment of LC3 is reduced heavily due to interaction with surface metalloprotease GP63^11^.

LC3 associated phagocytosis (LAP) is a kind of non-canonical autophagy which the parasite targets to manipulate host autophagic machinery accordingly. LC3 tagging on the surface of extending membrane is a crucial step and parasite alters it, thus acquiring larger entry to host cells but persisting to survive by inactivating canonical autophagic ways. The machinery is tempered by parasite attacking or residing cells leading to less production and tags of LC3 on membrane which in turn avoid maturation of autophagosome and fusion with lysosomes.

## 2. Methodology

### 2.1 Reconstruction of Autophagic Network

The complete autophagic network was analysed using the network analyser Cytoscape v.2.8.3^12^. The information of the proteins was retrieved from STRING v.10.5 and Gene Ontology Consortium databases; the proteins thus collected are based on their biochemical function and connectivity to the other proteins of autophagic machinery in *Homo sapiens* and *Mus musculus*. The total number of nodes present in the network is 100, which is a collection of autophagy related gene product proteins as well as transcription factors, cytokines and target products of autophagic machinery. From the given proteins and molecules, analysis is performed based on high values of clustering coefficient, closeness centrality, betweenness centrality, average shortest path length, outdegree and indegree.

*Clustering Coefficient* is a measure of the degree to which nodes in the network tend to cluster together, thus more the clustering coefficient is denser is network forming a closely-knit pattern. For a particular node, high clustering coefficient specifies its highest degree of connectivity. *Closeness Centrality* measures the mean distance from a node to other nodes, which means the more central a node is the closer it is to all other nodes. *Betweenness Centrality* measures the extent to which a node lies on paths between other nodes. Nodes with high betweenness may have considerable influence within a network as they control information passing between others. They would also affect the network and all nodes most when removed because they lie on the largest number of paths taken by messages. *Average Shortest Path Length* is a concept in network analysis that defines the average number of steps along the shortest paths for all possible pairs of network nodes. It measures the efficiency of information transport in a network. *Indegree* and *Outdegree* measure the connection of a particular node with incoming signals and outgoing signals respectively, to carry out the functions in the given network^13^.

### 2.2 Phylogenetic Tree Construction and Molecular Clock Analysis

*Phylogenetic tree construction:* Coding sequences of ATGs (ATG1-32) were retrieved from NCBI databases of all the organisms available ranging from prokaryotes to mammals. Multiple sequence alignment was performed using online multiple sequence alignment tool MultAlin. The fasta files obtained were converted to nexus format for phylogenetic tree construction. Phylogenetic tree was constructed using the software MrBayes v.3.2.^14^. MrBayes performs Bayesian inference of phylogeny using a variant of Markov chain Monte Carlo (MCMC). MrBayes uses MCMC methods to estimate the posterior distribution of model parameters. The posterior distribution is an average between knowledge about the parameters before data is observed (which is represented by the prior distribution) and the information about the parameters contained in the observed data (which is represented by the likelihood function). It provides a natural and principled way of combining prior information with data^14^. All inferences logically follow from Bayes’ theorem. MCMC, along with other numerical methods, makes computations tractable for virtually all parametric models. The nucleotide substitution model used during the tree construction was the GTR (General time reversible) model with gamma distributed rate variation across sites and a proportion of invariable sites.

MCMC simulation was run for 1 lakh to 40 lakh generations, depending on standard deviation of split frequency of each ATG, trees were sampled every 1000 generations. Analysis was run until a standard deviation of split frequencies were below 0.01. Parameters and corresponding trees were summarized and used later for analysis. Phylogenetic tree of all the ATGs was visualized using FigTree v.1.4.3^15^

*Molecular clock analysis:* Information regarding the out-groups was obtained using the phylogenetic tree construction. Only those ATGs which play utmost important role in machinery is taken for study which includes ATG1, ATG2, ATG4, ATG5, ATG7, ATG8, ATG9, ATG10, ATG12, ATG16 and ATG18. Relaxed molecular clock was constructed using MrBayes v.3.2.1.

MCMC simulation was run for one lakh to 10 lakh generations, again depending on each ATG. The model used for constructing the molecular clock was the independent gamma rates model (IGR model). Molecular clocks of aforementioned ATGs were constructed and visualized using the FigTree v.1.4.3.

### 2.3 Reconstruction of Autophagic Model by Mathematical Modelling

#### 2.3.1 Model Reconstruction and Simulation

Signaling network reconstruction is the integration of information that describes the biochemical changes that occur in a specified network. It helps gaining deeper understanding of interactions between the signaling proteins with themselves and surrounding as well as explaining the complexity of biological system at higher level. Quantitative modeling of these interactions plays an important role in understanding fundamental intra- and inter-cellular processes^3^.

The interactions between the signaling components were entered as elemental chemical reactions in MATLAB v.7.11.1.866 SimBiology toolbox, which uses the Systems Biology Markup Language (SBML) machine language. The kinetic rate laws (mass action for association and dissociation reactions, translocation and conversion, Henri–Michaelis– Menten for phosphorylation/ dephosphorylation/ ubiquitination, Hills equation for gene expression) and initial concentrations associated with the reactions were also defined in the toolbox, which were applied for a desired behaviour.

The reconstructed network was numerically simulated using Stiff Deterministic ODE15s solver (SimBiology toolbox) which generates the first order nonlinear ODEs for each node, thus defining the mathematical structure of the model. Complex biological systems, such as the signaling network can be viewed as networks of chemical reactions that can be analysed mathematically using ODEs, which is the most common simulation approach used in computational systems biology. ODE helps in determining time dependent changes i.e. the time series data of the concentrations of the signaling proteins and protein complexes and thus the dynamics associated with it. The model was exported to Copasi v.4.8.35^16^ as a SBML file to generate the time series data.

#### 2.3.2 Parameter Estimation

The equations outlining the chemical transformation in the signaling flow use variables as concentration of protein. Often these parameters are unknown, difficult and hard to measure. In such situation parameter estimation is done i.e. the unknown parameters are determined indirectly using computational biology tools. The parameters for reconstructed signaling network were manually trained within a range of parameter range that was further tuned until the model was simulated to obtain a desired output with algorithmic data applicable due to lack of homogeneity in experimental measurements^3^.

#### 2.3.4 Sensitivity Analysis

It plays an important role in dynamic analysis of systems biology models. Sensitivity analysis is used to determine how sensitive a model is to variables changes in the model, which helps study the randomness associated with the estimated parameters and thus build assurance in the model^3^.

It was calculated using the SimBiology toolbox, Sensitivity Analysis option that calculates the sensitivity coefficients by combining the original ODE system for a model with the auxiliary differential equations for the sensitivities. The additional equations are derivatives of the original equations with respect to parameters. It calculates the time-dependent sensitivities of all the species with respect to species initial conditions and parameter values in the model.

In this paper, signaling network of two different cases has been taken into consideration:

***CASE:1 Interplay between TNFα and IL10 for autophagy in macrophages in leishmaniasis***-In this case, two models were developed in which one was for induction of autophagy by TNFα and the second model was constructed based on inhibition of autophagy by IL10 signaling pathways. Moreover, a model for general study of LAP was also reconstructed.

***CASE:2 Interaction between canonical and non-canonical autophagy in leishmaniasis****-* In this case, three models were developed in which one explains the signaling pathway compiling both canonical and non-canonical autophagy, second model is designed that explains only canonical autophagic pathway excluding LAP and the last model is reconstructed to explain only non-canonical autophagy which is LAP excluding canonical autophagy.

### 2.4 Principal Component Analysis and Model Reduction

#### 2.4.1 Principal Component Analysis (PCA)

Generation of a new set of variables (principal component score), called principal components, from the multivariate data sets is done using MATLAB v.7.11.1.866. Each principal component is a linear combination of the original variables. All the principal components are orthogonal to each other, so there is no redundant information. PCA was done in MATLAB using the function i.e. [score_coefficient] = princomp (a) where, “a” is the y matrix of sensitivity coefficients of each species in the signaling network. This command returns principal component coefficient and score of the sensitivity matrix, used to identify the principal component i.e. significant individual reactions in the signaling network. The graphical representation of the derived score coefficient is visualised using SigmaPlot v.10.0 by scatter plot formation^3^.

#### 2.4.2 Model Reduction

Model reduction is a systematic method for eliminating reactions that do not contribute considerably to network output. In this study flux analysis, sensitivity analysis and concentration at which the species shows the highest sensitivity was combined to reduce the dimensionality of the model. The concentrations were retrieved from the time series data using Copasi v.4.8.35. The graphical representation of the collected values is visualised using SigmaPlot v.10.0 by 3D-mesh plot formation^3^.

### 2.5 Flux Analysis

Flux analysis is applicable for systems that are in a pseudo-steady state. Under this condition, the differential equation system of metabolite mass balances reduces to a linear equation system, which relies solely on the known stoichiometry of the biochemical reaction network. For this system flux analysis for each reaction is done in Copasi v.4.8.35 under the task - Steady State Analysis and graphical representation is created using MS Excel v.2010^3^.

## 3. Results and Discussion

### 3.1 Autophagic Network Analysis

From the analysis of network of autophagy in human and mouse, the crucial crosstalk point molecules obtained are Beclin1, ATG9, ULK and PI3P (Fig 3). From this result, we can understand that these molecules play a critical role in signaling machinery of autophagy i.e. they could be used as target to perturb the network accordingly and help in decrease of parasite load.

**Fig. 3:**
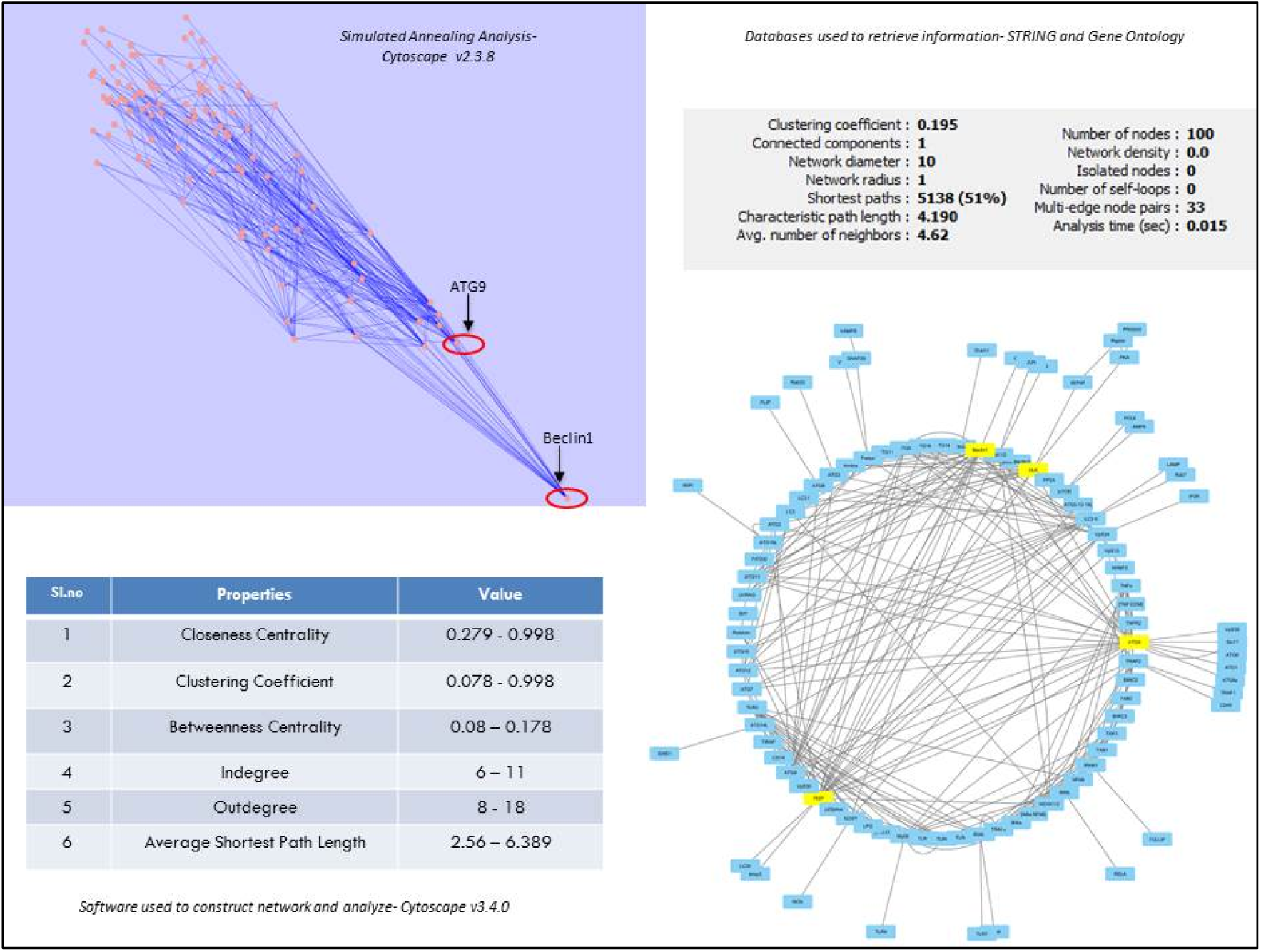
Autophagy network analysis and (table 4.1) parameters used to get crucial crosstalk points of network.

**Fig. 4:**
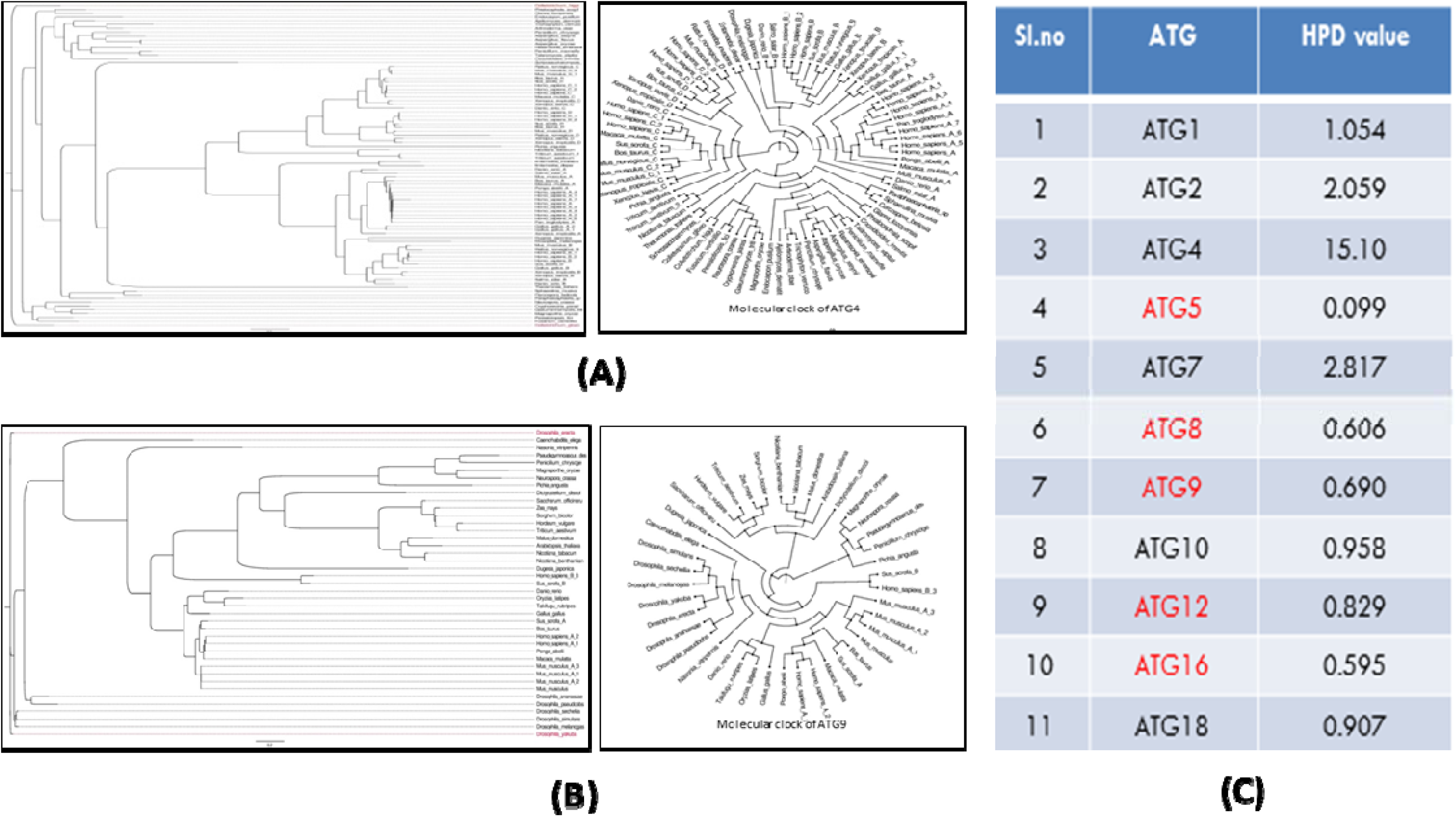
Phylogenetic and molecular clock: (A) ATG 4 (B) ATG 9 (C) Table showing 95% HPD value of various ATG.

### 3.2 Phylogenetic Tree Construction and Molecular Clock Analysis

Due to presence of core autophagic proteins i.e. autophagy related gene products in the crucial components; they are taken for phylogenetic analysis. From all the coding sequences of different autophagy related genes, few genes are further processed in molecular clock that are defined well in the process of autophagy. Moreover, from this analysis, we come to refine our result as ATG5, ATG8, ATG9, ATG12 and ATG16 are conserved amongst variable taxon and could be used further to construct mathematical model to analyse principal components of autophagic machinery (Fig.4).

### 3.3 Mathematical Modelling, Principal Component Analysis and Model Reduction

With the mentioned concentration, reaction laws and parameters models for two different set of cases were constructed.

#### Case 1- <u>Interplay between TNFα and IL10 for autophagy in macrophages in leishmaniasis:</u>

##### **A.** For Induction of Autophagy

##### B. For Inhibtion of Autophagy

In this case of interplay between two cytokines, the induction (Fig 5A) and inhibition (Fig 5B) of autophagy is studied where TNFα is determined to be the cause of induction and IL10 for inhibition. In the model system concerning the induction of autophagy, the principal components ATG9 and PI3P are obtained, whereas, for inhibition phosphorylated AKT in membrane and phosphorylated mTOR generated by AKT pathway are obtained. Model is then further reduced to two reactions viz., interaction between ATG9 and PI3P as well as phosphorylation of mTOR.

**Fig. 5.**
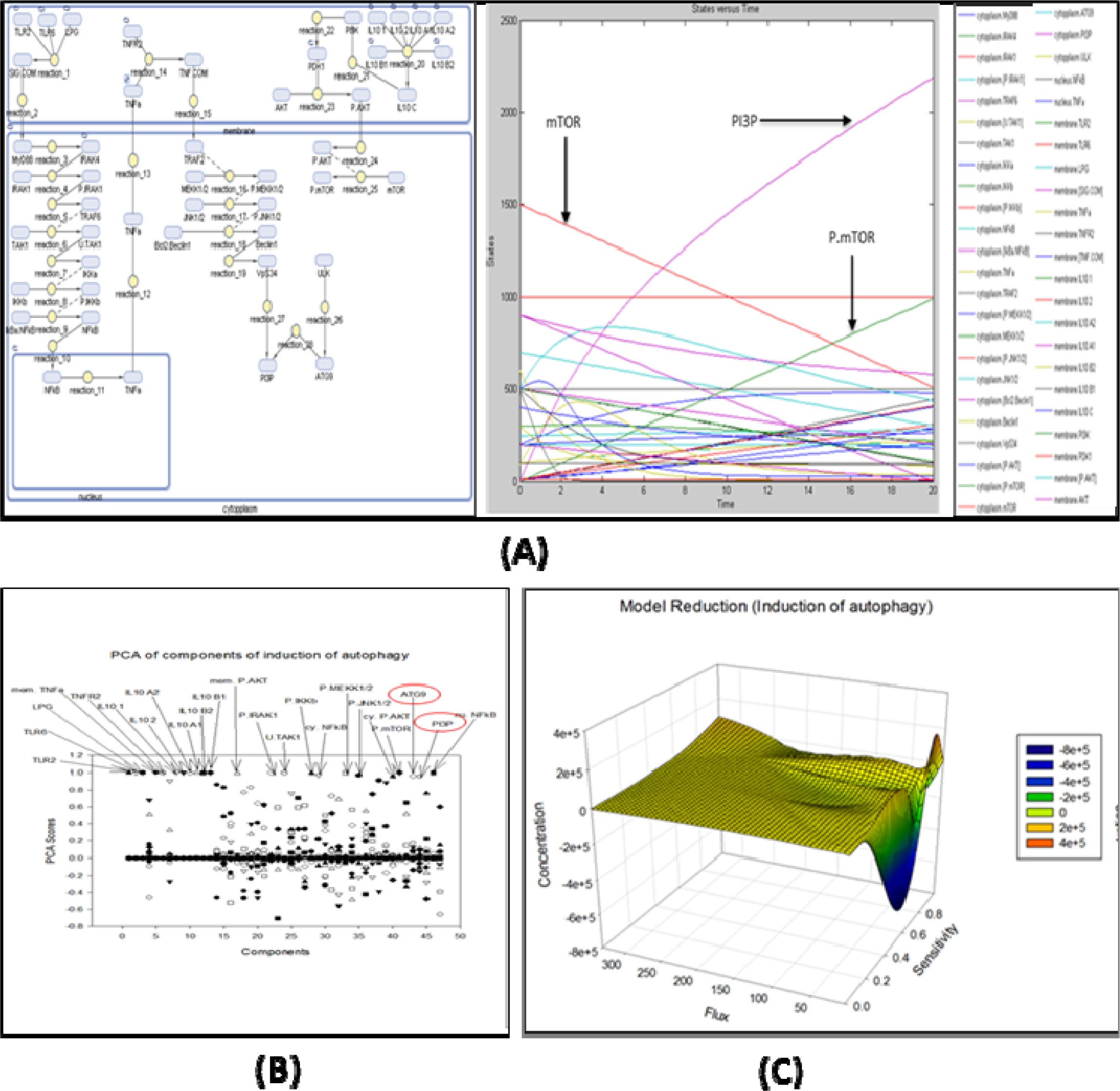
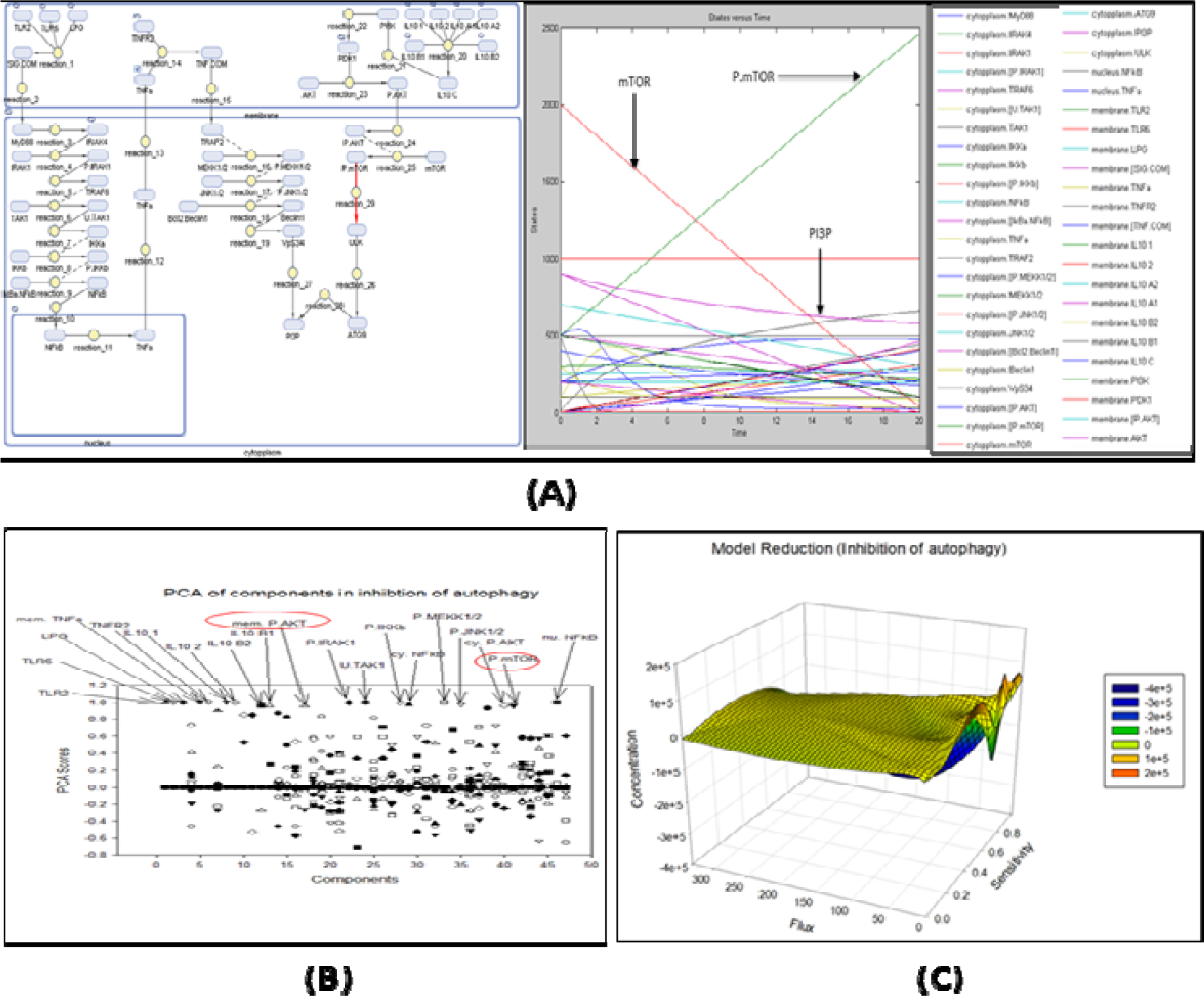
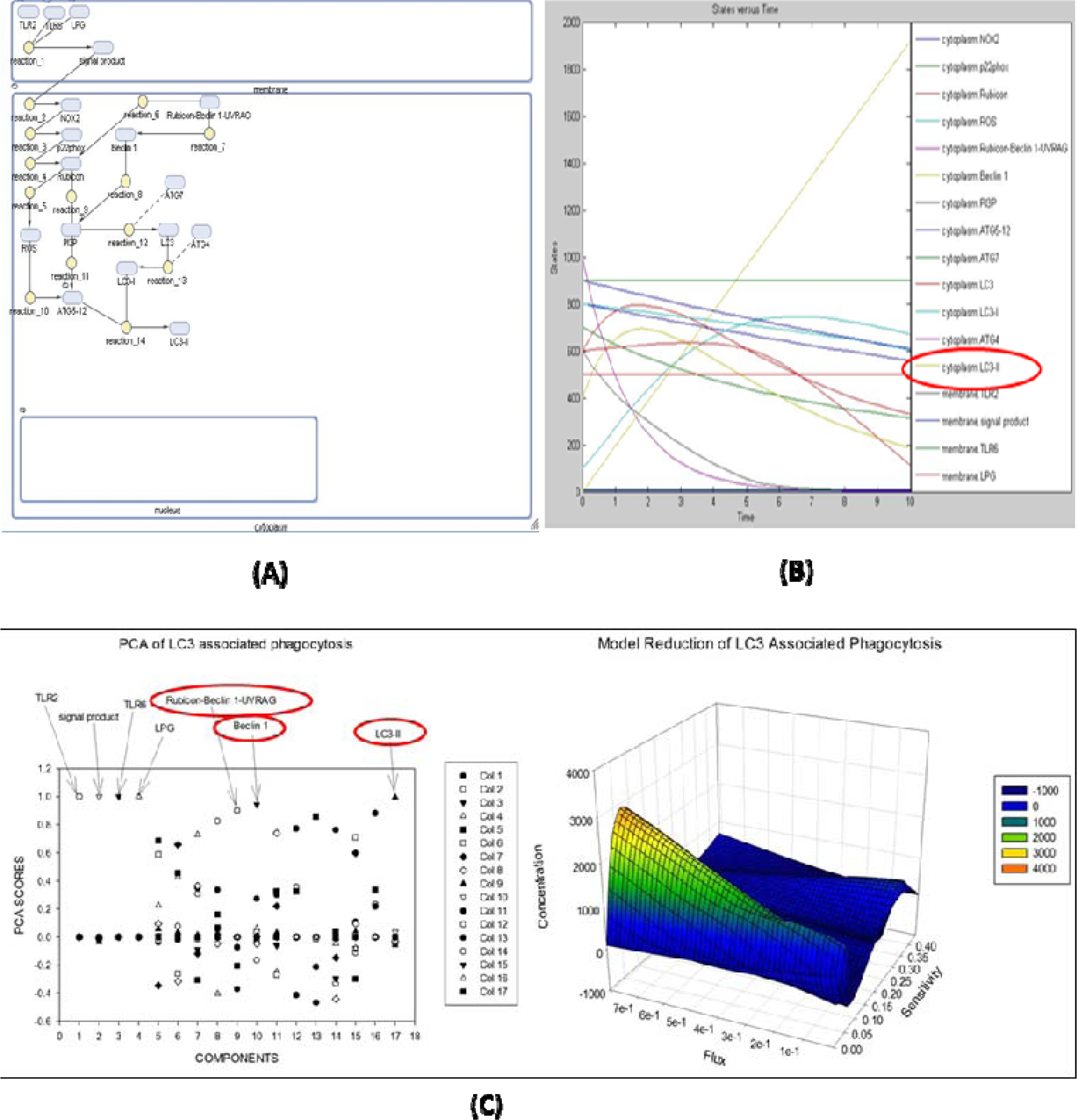
A: - Interplay between TNFα and IL10 for autophagy in macrophages in leishmaniasis: (A) Model reconstruction of induction of autophagy and simulated graph (B) Principle component analysis showing mTOR and PI3P as an important component (c) Graphical representation of model reduction. B: - Interplay between TNFα and IL10 for autophagy in macrophages in leishmaniasis: (A) Mode reconstruction of inhibition of autophagy and simulated graph (B) Principle component analysis showing mTOR and membrane phosphorylated AKT as an important component (c) Graphical representation of model reduction. C: (A) Model reconstruction of LC3 associated phagocytosis and its simulated graph constructed through Simbiology (B) Principle component analysis showing Beclin1, Rubicon and LC3 as an important component (C) Graphical representation of model reduction.

##### **C.** Model reconstruction for LAP (LC3 associated phagocytosis)-

In case of general study on LAP, Rubicon-Beclin1-UVRAG complex, Beclin1 and LC3II are produced as principal component (Fig 5C).

#### Case 2-Interaction between canonical and non-canonical autophagy in leishmaniasis

##### A. Model reconstruction of system having combination of canonical and non-canonical autophagy

##### B. Model reconstruction of system having canonical autophagy signaling excluding LAP

##### C. Model reconstruction of system having only non-canonical autophagy (LAP) excluding canonical autophagy

In the case of interaction between canonical and non-canonical autophagy in leishmaniasis (Fig 6A,6B and 6C), the principal components in the system obtained are p22phox, LC3 variants, ATG4, ULK and ROS (for combined study of canonical and non-canonical autophagy), TRAF2, ULK, ATG4, LC3, ATG5-12-16L complex and Beclin1 (for system with only canonical autophagic machinery signaling) and p22phox, LC3 variants, ATG4 and ROS (for system with only non-canonical autophagic machinery signaling). The reduced model elucidates four important cross-talk point molecules as ATG4, ATG9, Beclin1 and Rubicon.

**Fig. 6.**
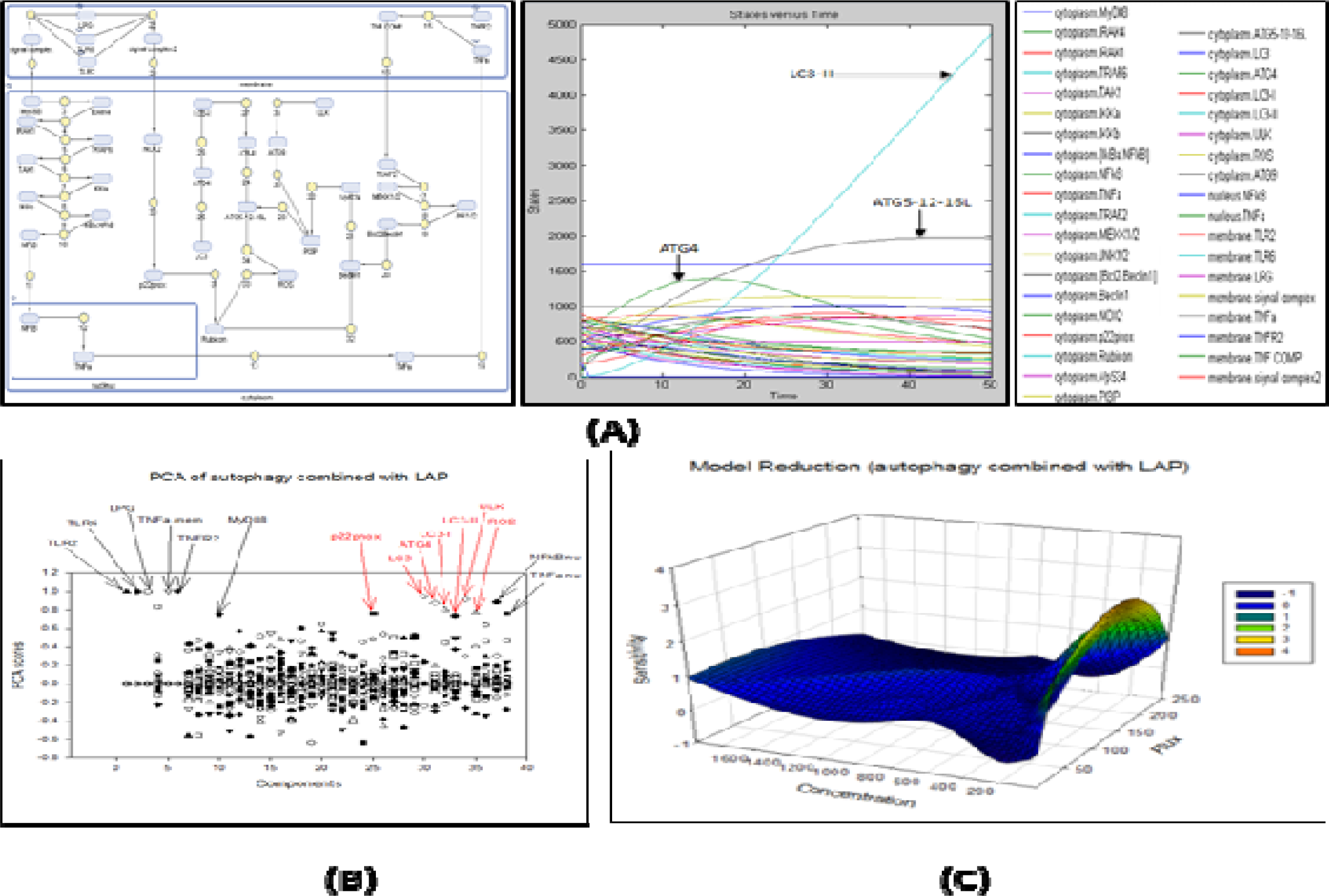
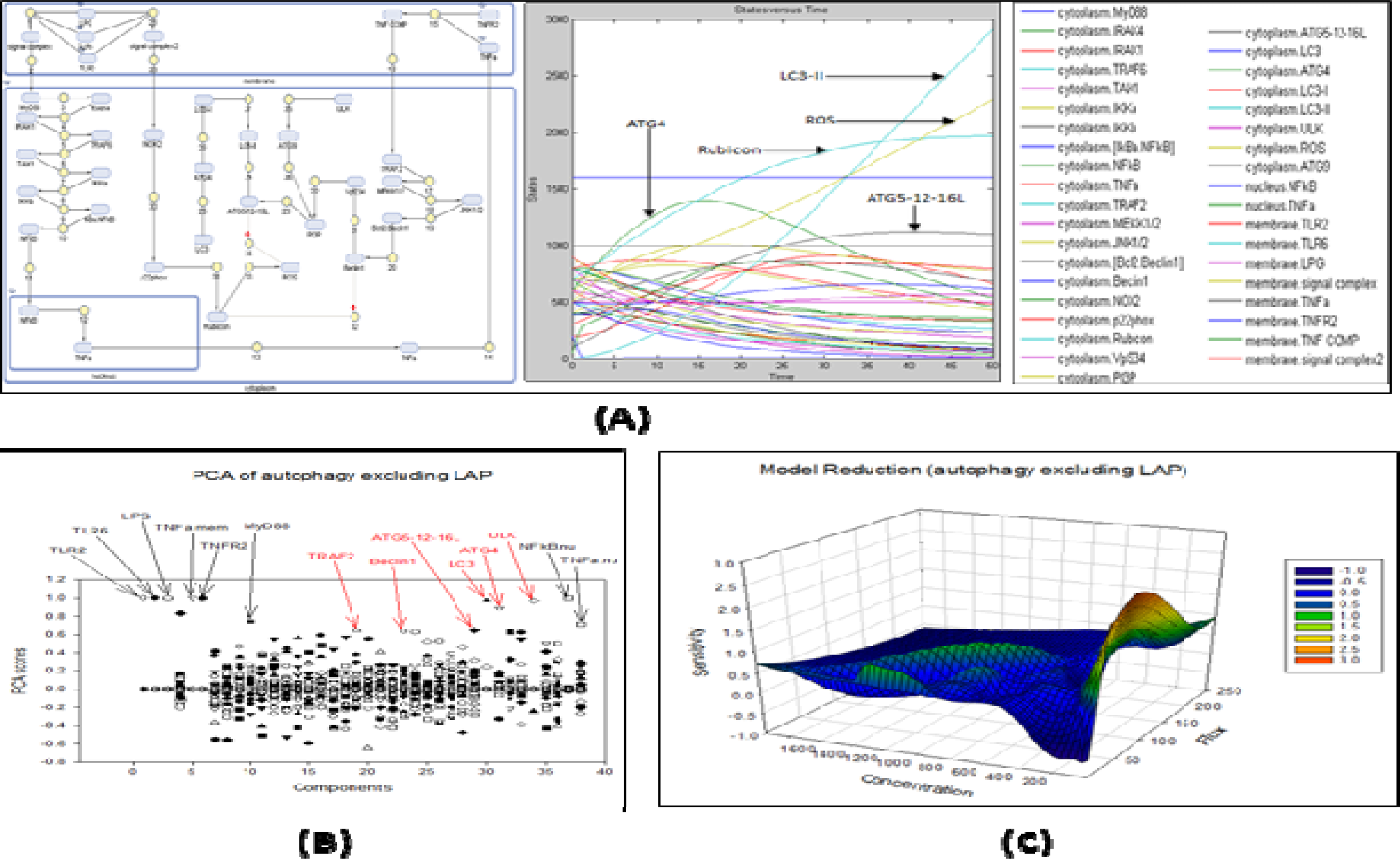
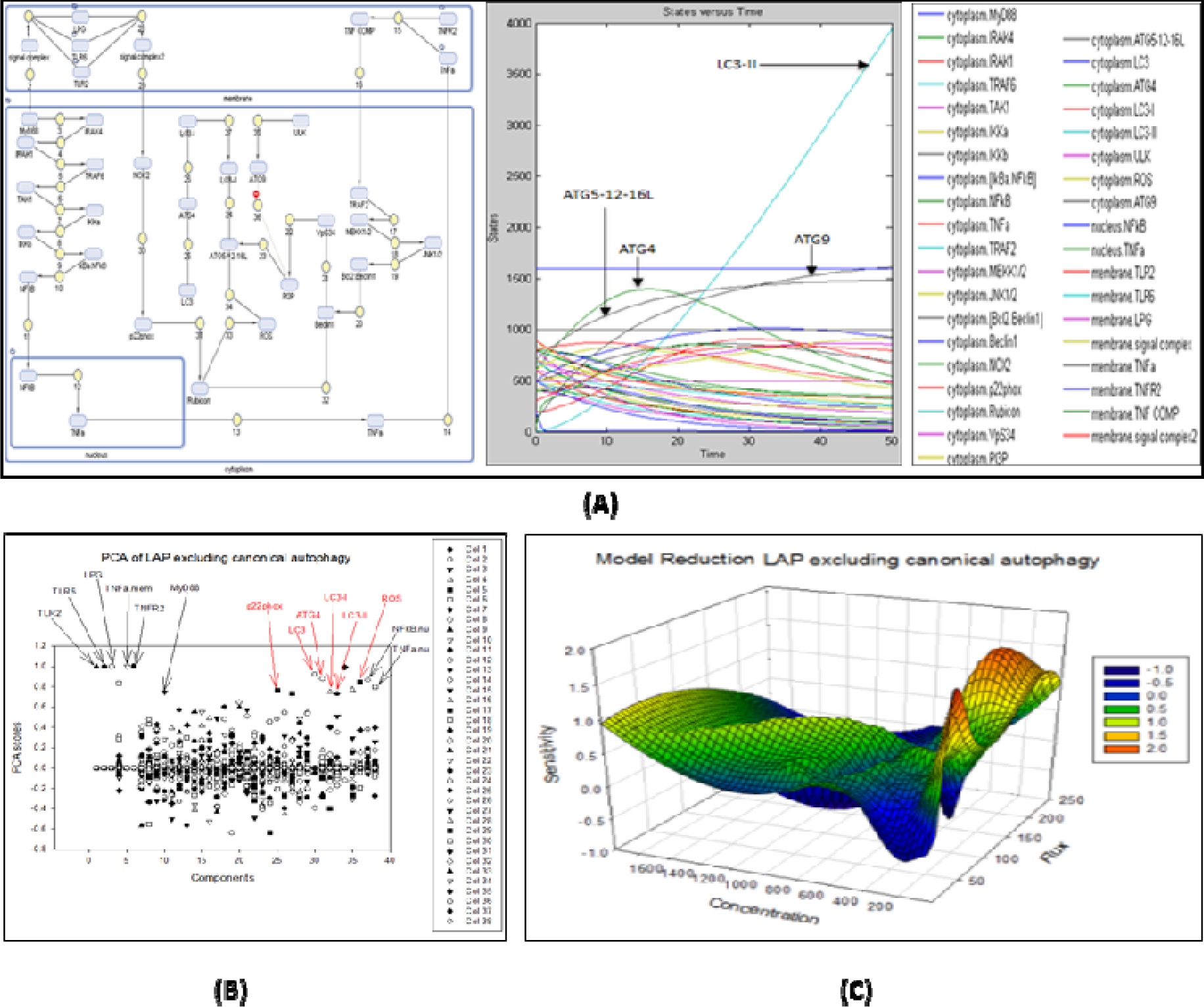
A-Interaction between canonical and non-canonical autophagy in leishmaniasis: (Model reconstruction of system having combination of canonical and non-canonical autophagy and its simulated graph (B) Principle component analysis depicting ATG4 as an important component (C) Graphical representation of model reduction. B-Interaction between canonical and non-canonical autophagy in leishmaniasis:(A) Model reconstruction of system having canonical autophagy signaling excluding LAP and its simulated graph (B) Principle component analysis showing ATG4, Beclin1 and LC3 as an important component (C) Graphical representation of model reduction. C-Interaction between canonical and non-canonical autophagy in leishmaniasis:(A) Model reconstruction of system having only non-canonical autophagy (LAP) excluding canonical autophagy and its simulated graph (B) Principle component analysis showing ATG4, Beclin1 and LC3 as an important component (C) Graphical representation of model reduction.

**Fig. 7:**
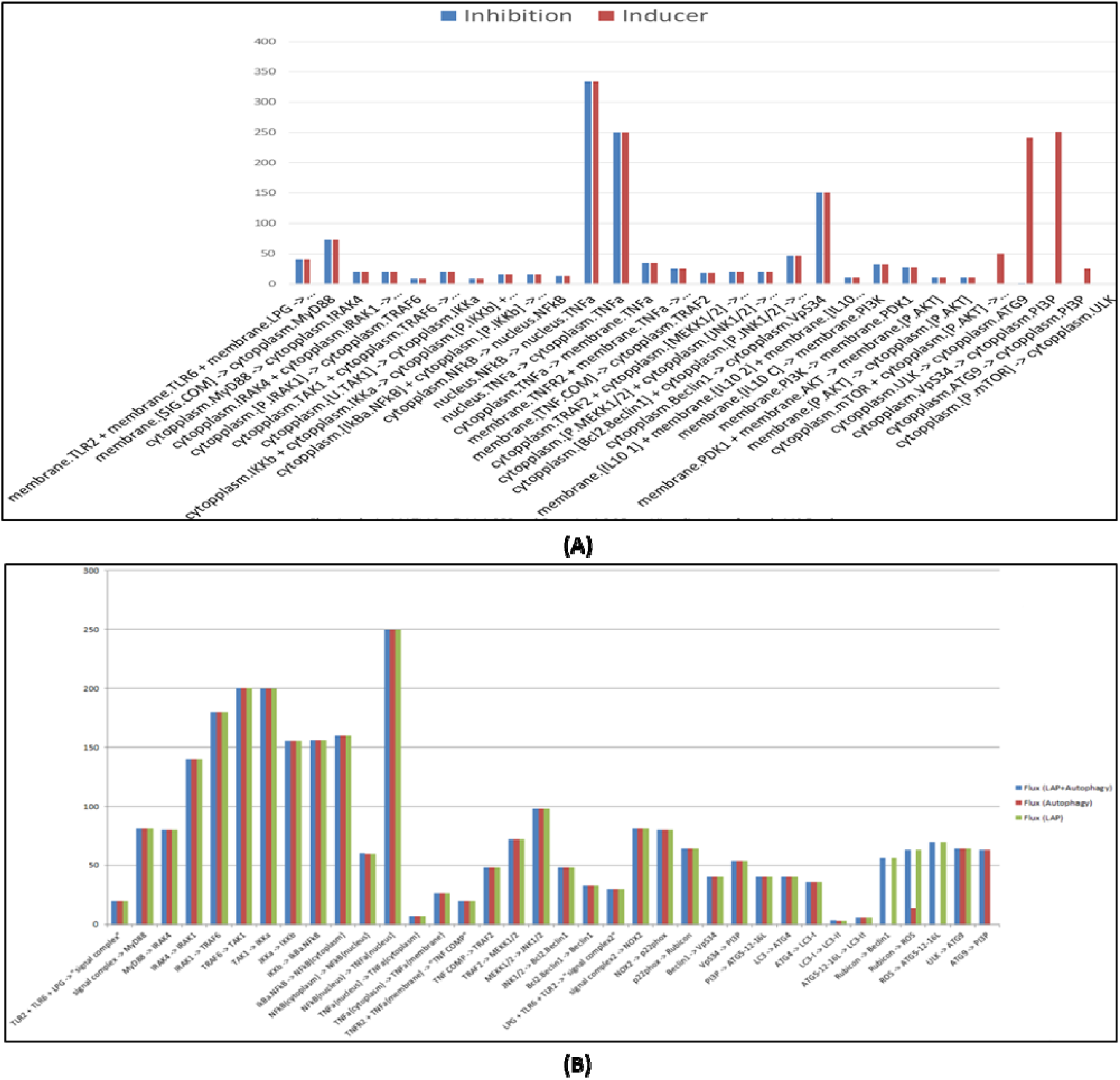
Flux analysis: (A) Showing interplay between TNFα and IL10 for autophagy in macrophages in leishmaniasis (B) Showing interaction between canonical and non-canonical autophagy in leishmaniasis.

### 3.4. Flux Analysis

The major reaction of the system is collection of LC3II, link of ATG9 and PI3P and disassembling of Beclin1 from the Rubicon complex in study of interplay between TNFα and IL10 governing autophagy. Whereas, in case explaining interaction between canonical and non-canonical autophagy in leishmaniasis four reactions are obtained as crucial reactions that govern the whole system. They are activity of ATG9, ATG4, activity of Beclin1 and processing and accumulation of LC3 on autophagosomal membrane.

## 4. Conclusion

During the course of evolution ATG9 remains conserved and is found to be an important component in autophagic machinery including Beclin 1. ATG9’s is observed to have a negative feedback role with interaction to PI3P in LC3 associated phagocytosis since there is low level of LC3-II accumulation when the non-canonical autophagic pathway is switched off. The simulated graph denotes significant change in concentration of LC3 accumulation in the system which helps PI3P accumulation to growing membrane but decreases affinity of LC3 attachment when LAP is not working in tandem. Study of leishmanial ATG9 may help understand the role of PI3P usage in parasite for nutrition and thus may help parasite prevalence in the host. The interaction between leishmanial ATG9 and PI3P shows its similarity with host autophagic machinery ATG9, hence it could be concluded that if the parasitic ATG9 is found to interfere with the host machinery, it could be targeted. Moreover, ATG9 is linked with lipid accumulation for autophagosome formation and interaction with PI3P which governs membrane dynamics of host and thus proves to be an important factor for parasite prevalence.

## 5. Future Perspectives

The interconnected biochemical pathways shed an important knowledge governing the core cellular processes about autophagy into therapeutics. Macroautophagy plays an important role in the inflammatory conditions, and more so, when infection with *L.major* is taken into account. Small molecule/inhibitor designed against the target identified (Atg9) could help the remedial measures not getting averse both for autophagic machinery and other pathophysiological processes associated with inflammatory diseases.

## 8. Acknowledgements

The authors would like to thank the Director, National Centre for Cell Science (NCCS) for supporting the Bioinformatics and High Performance Computing Facility (BHPCF) at NCCS, Pune, India.

